# Spectral distribution dynamics across different attentional priority states

**DOI:** 10.1101/2021.12.02.470964

**Authors:** Mattia Pietrelli, Jason Samaha, Bradley R. Postle

## Abstract

Anticipatory covert spatial attention improves performance on tests of visual detection and discrimination, and shifts are accompanied by decreases and increases of alpha-band power at EEG electrodes corresponding to the attended and unattended location, respectively. Although the increase at the unattended location is often interpreted as an active mechanism (e.g., inhibiting processing at the unattended location), most experiments cannot rule out the alternative possibility that it is a secondary consequence of selection elsewhere. To adjudicate between these accounts, we designed a Posner-style cuing task in which male and female human participants made orientation judgments of targets appearing at one of four locations: *up, down, right*, or *left*. Critically, trials were blocked such that within a block the locations along one meridian alternated in status between *attended* and *unattended*, and targets never appeared at the other two, making them *irrelevant*. Analyses of the concurrently measured EEG signal were carried out on traditional narrowband alpha (8-14 Hz), as well as on two components resulting from the decomposition of this signal: periodic alpha; and the slope of the aperiodic 1/f-like component. Although data from *right*-*left* blocks replicated the familiar pattern of lateralized asymmetry in narrowband alpha power, with neither alpha signal could we find evidence for any difference in the time course at *unattended* versus *irrelevant* locations, an outcome consistent with the secondary-consequence interpretation of attention-related dynamics in the alpha band. Additionally, 1/f slope was lower at *attended* and *unattended* locations, relative to *irrelevant*, suggesting a tonic adjustment of physiological state.

## Introduction

Valid expectations about where a visual stimulus will appear improve performance on tests of visual detection and discrimination (Posner, 1980). With electroencephalography (EEG) the onset of a spatial cue triggers a decrease in the power of oscillations in the alpha band (∼8-14 Hz; henceforth “alpha”) at posterior electrodes contralateral to the cued location, and a commensurate increase in alpha power at electrodes ipsilateral to the cued location (Thut et al., 2006; Rihs et al., 2009; Capilla et al., 2014). However, consensus is lacking about the functional interpretation of this familiar pattern: Are these attention-related changes in alpha dynamics reflective of active mechanisms that implement attentional control (Sadaghiani and Kleinschmidt, 2016), or are they byproducts of attention-related state changes that were caused by other mechanisms (i.e., a “secondary consequence”; Antonov et al., 2020)? As an example of an active mechanism, one influential account holds that the increase in power at “unattended electrodes” reflects the active inhibition of processing at that location, presumably to suppress the processing of distracting information (Kelly et al., 2006; Jensen and Mazaheri, 2010). Alternatively, a “secondary-consequence” account interprets this same effect as a consequence of the withdrawal of attention (Foster and Awh, 2019; Antonov et al., 2020). An analogy for the secondary-consequence account is eye closing, for which the increase in power of posterior alpha is a consequence of the loss of visual input, not the operation of a mechanism that caused eyelid closure.

To adjudicate between these accounts, we modified the standard spatial-cuing procedure – which compares *attended* vs. *unattended* electrodes -- to create a third class of electrodes: *irrelevant* electrodes corresponding to locations that would never be cued during a block of trials. On any block of trials, targets would only appear to the *left* or *right* of fixation, but never *above* or *below* fixation, thereby making *above* and *below irrelevant* during this block of trials. If the increase in alpha power at the *unattended* location reflects an active mechanism, it should not be observed at the *irrelevant* locations. (E.g., in the motor control, memory and working memory literature, focal increases in alpha power are found when top-down inhibitory control is needed to withhold or otherwise control the execution of a response to an otherwise prepotent location (Klimesch et al., 2007; Jensen and Mazaheri, 2010). *Irrelevant* areas should not have such prepotency.) If, alternatively, the increase in alpha power at the *unattended* location is a secondary consequence of the selection of a different location, *irrelevant* locations might also experience a return to a baseline state.

One potential concern about our design is that one of the two accounts – secondary consequence -- predicts the absence of evidence for a statistical difference between *unattended* and *irrelevant* alpha power. Therefore, were this to be the outcome, we planned to assess whether some other component in the same data does discriminate *unattended* from *irrelevant*. More specifically, we planned to decompose the EEG data into periodic and aperiodic components. Although task-related modulation in a predefined frequency band (e.g., alpha) is most often interpreted as a change in the power of an oscillator, at least some of that change might be the result any of several other factors, including a change of the center of frequency of an oscillator (Haegens et al., 2014; Mierau et al., 2017), a change of the bandwidth of an oscillator, or a change in the 1/f-like aperiodic component on which the oscillator sits (Voytek et al., 2015; Donoghue et al., 2020a). Because these factors are mathematically independent, it is possible that the neural generators underlying them may also vary independently. Consequently, if “traditional alpha,” generated with bandpass-filtering, failed to discriminate *unattended* locations from *irrelevant* ones, we reasoned that perhaps one of these decomposed components of the EEG signal would be successful.

## Materials and methods

### Subjects

Nineteen healthy subjects (8 males and 11 females; age M = 22.6, S = 4.7) were recruited from the University of Wisconsin–Madison community and were compensated monetary. This study sample size was based on previous studies that have influenced our thinking about attention-related alpha dynamics (Thut et al., 2006; Rihs et al., 2009), in order to facilitate contextualization of our findings with them. All subjects reported right-handedness and normal or corrected-to-normal visual acuity and color vision. None reported any medical, neurological, or psychiatric disorder. The University of Wisconsin–Madison Health Sciences Institutional Review Board approved the study.

### Experimental design

The task was two-alternative forced-choice discrimination of orientation (clockwise/counterclockwise), with a symbolic cue preceding the target and carrying information about where and when the target would appear. Trials were administered in alternating 70-trial blocks in which target stimuli appeared exclusively at a location 7.5 degrees of visual angle (DVA) to the left or to the right of central fixation, or exclusively 7.5 DVA above or below fixation. Overall, subjects completed 5 horizontal and 5 vertical blocks, for a total of 700 trials per subject.

#### Experimental procedure

Subjects completed the experiment in one session, while seated in an acoustically shielded and dimly lit room. Stimulus presentation was controlled with MatLab (2020b, The MathWorks, Natick, MA) and presented on a 53×30cm screen (refresh rate, 60 Hz) at a viewing distance of ∼57 cm. Throughout the experiment the background screen was gray. Prior to the 10 experimental blocks, each subject first completed a titration procedure to determine presentation parameters that produced discrimination performance near 75% accuracy, then a training block using those parameters. For clarity, the titration procedure and training block will be described after the description of the main experiment.

In the main experiment, each trial started with the 200 ms presentation of a symbolic cue (an arrowhead) in the center of the screen. The direction of the arrow indicated the location of the forthcoming target with 75% validity; on the remaining 25% of trials the target appeared in the opposite location on the same meridian. The color of the arrow contained information about the cue-to-target interval (CTI): a magenta arrow indicated, with 100% validity, a CTI of 650 ms, whereas a green arrow indicated that the CTI could come from one of four durations (650 ms, 900 ms, 1150 ms, 1400 ms) drawn randomly from an exponential distribution. From the full set of 700 trials, only the 400 trials trials with a CTI of 650 ms were used for the analyses reported here.

The cue was replaced by a white fixation cross (2°), then the target (50 ms), then a backward mask (50 ms). The target was a Gabor patch, 80×80 pixels (approximately 2°) in size, with spatial frequency of one cycle every 10 pixels, oriented either 45° clockwise or counterclockwise of vertical. Novel masks were generated for each trial by filling in a circular aperture the size of Gabor with randomly arranged black and white pixels. Subjects were instructed to make a clockwise/counterclockwise orientation discrimination of each target, and to register their decision as quickly and accurately as possible by pressing the left arrow key on a keyboard for a counterclockwise orientation (index finger of right hand) and the right arrow key for a clockwise orientation (middle finger of right hand). After each orientation response, subjects then rated the subjective visibility of the target, using the Perceptual Awareness Scale (PAS) rating (Ramsøy and Overgaard, 2004): The sentence “Perceptual awareness rating?” was presented, along with the digits “1” “2” “3” “4”, and subjects indicated their rating by selecting the corresponding key on the keyboard with their left hand. Subjects were instructed to press “1” when they had no subjective experience of seeing the target; to press “2” when they perceived a distinction between target and mask, but did not perceive the target’s orientation; to press “3” when they had an impression that they had perceived the target, but that it was not clearly visible; and to press 4 when they were confident that they had seen the target clearly. For this, subjects were asked to “think carefully” about their visibility rating, and to take as much time as they needed before responding. In order validate this aspect of the procedure, no target was present on 20% of trials (just a black 80×80 patch without any stripe for 50 ms, followed by the 50 ms backward mask).

Inferential statistical analyses on the behavioral data were performed using STATISTICA (Version 12.0; StatSoft), and post-hoc comparisons were conducted using Newman–Keuls tests.

#### Titration procedure

Prior to the main experiment, each subject completed two runs through a titration procedure, one each for vertical and for horizontal locations, during which the contrast level of the target was varied (via the QUEST procedure) to obtain levels of orientation discrimination of 75% correct (Watson and Pelli, 1983; King-Smith et al., 1994). Each trial of the titration runs was identical to the experimental blocks, except no symbolic cue was presented, no visibility report was requested, and feedback was provided after each response.

#### Training procedure

After titration, subjects received instruction via progressive introduction (from the trial structure of the titration procedure) to elements from the experimental task, using the contrast level previously found during the titration procedure. First, they were instructed about the meaning of the direction of the cue and performed 20 trials with a white cue. Next, were instructed about the meaning of the color of the cue and performed 20 additional trials with the cue containing both the location and timing information. Finally, they were instructed on how to respond on the PAS, and during 40 remaining trials the visibility rating screen was presented after each orientation response.

### EEG recording and preprocessing

EEG signal was recorded from 60 Ag/AgCl electrodes with positions conforming to the extended 10-20 international system. Recordings were made using a forehead reference electrode and amplified with an Eximia 60-channels amplifier (Nextim; Helsinki, Finland) with a sampling rate of 1450 Hz. Preprocessing and analysis were conducted with MatLab (2020b, The MathWorks, Natick, MA) using custom routines and EEGLAB (v. 2019/1) toolbox (Delorme and Makeig, 2004). Data were down-sampled offline to 1000 Hz and were divided into epochs ranging between -1000 to +1400 ms around cue onset. Epochs with muscle artifacts were identified by a trained operator and discarded (M = 2.3%, STD = 4.0%). After reducing data dimensionality to 32 components based on the Principal Component Analysis, horizontal and vertical eye-movement artifacts were visually identified and removed by means of Independent Component Analysis (ICA) by a trained operator (M = 1.0%, STD = 0.7%).

#### Time-frequency and periodic/aperiodic decomposition

From the cleaned data, a complex Morlet wavelet decomposition (Cohen, 2019) was applied to each single epoch (1-50 Hz, 0.35 Hz frequencies bins, 3 to 10 cycles) with a custom routine in the Fieldtrip toolbox environment (Oostenveld et al., 2011; version 2021/03/11). The resultant time-frequency series were down-sampled from 1800 (1000 Hz) to 200 (111 Hz) timepoints, to reduce processing time for subsequent analyzes. In order to track alpha power activity across the different attentional priority conditions, power values were averaged across alpha band interval 8-14 Hz (Thut et al., 2006; Rihs et al., 2009) and baseline corrected (by subtraction) to the average across all trials of the pre-cue period between -500 and 0 ms. It is on these data, from here forward “narrowband alpha power,” that we carried out the analyses intended to adjudicate between inhibition vs. idling accounts of alpha dynamics during spatial cuing.

After analyzing the narrowband alpha power data, we decomposed the EEG data into putatively oscillatory and aperiodic components by carrying out a parametrization of the spectral distribution with the “fitting-oscillations-and-one-over-f” (FOOOF) toolbox (Donoghue et al., 2020b). For each subject, a FOOOF routine was run on the time-frequency series, estimating the aperiodic-adjusted power at the peak of alpha and the slope of the 1/f aperiodic component of the spectral distribution, separately for each electrode, each trial, and each timepoint. Finally, the estimated aperiodic-adjusted power at the peak of alpha and the slope of the 1/f aperiodic component values were baseline corrected (by subtraction) to the across-trial average of the pre-cue period between -500 and 0 ms, to produce what we will refer to as “decomposed alpha power” and “1/f slope.”

### Electrode selection

The main challenge of our procedure was the need to isolate signals selective for each of the four target locations. Indeed, for the selection of “up” and “down” locations on the vertical meridian there is not a procedure analogous to selecting a stereotypical set of contralateral electrodes as is typically done for tasks using only left and right locations. Moreover, the number of electrodes selected per ROI is often arbitrary, even for typical left/right locations. Consequently, for all of our hypothesis tests we employed L1-regularized logistic regression classifiers (Tibshirani, 1996; from now will be called just LASSO) to select the electrodes that would make up the region of interest (ROI) that preferentially represented each of the four tested locations.

#### A priori ROIs

Prior to hypothesis testing, to confirm that our data replicated established findings, we constructed a priori ROIs to isolate signal corresponding to left (P8, P10 and O2) and right (P7, P9 and O1) target locations (Thut et al., 2006; Rihs et al., 2009). Fig. 1. illustrates that, with these a priori ROIs, data from horizontal blocks showed the expected pattern of greater narrowband alpha power at electrodes corresponding to unattended than attended locations. Visual inspection of the data from Fig. 1 was also used to select the time interval of 1000-1200 ms with which to train classifiers to identify the electrodes for construction of *up, right, down*, and *left* ROIs.

**Figure 1.**
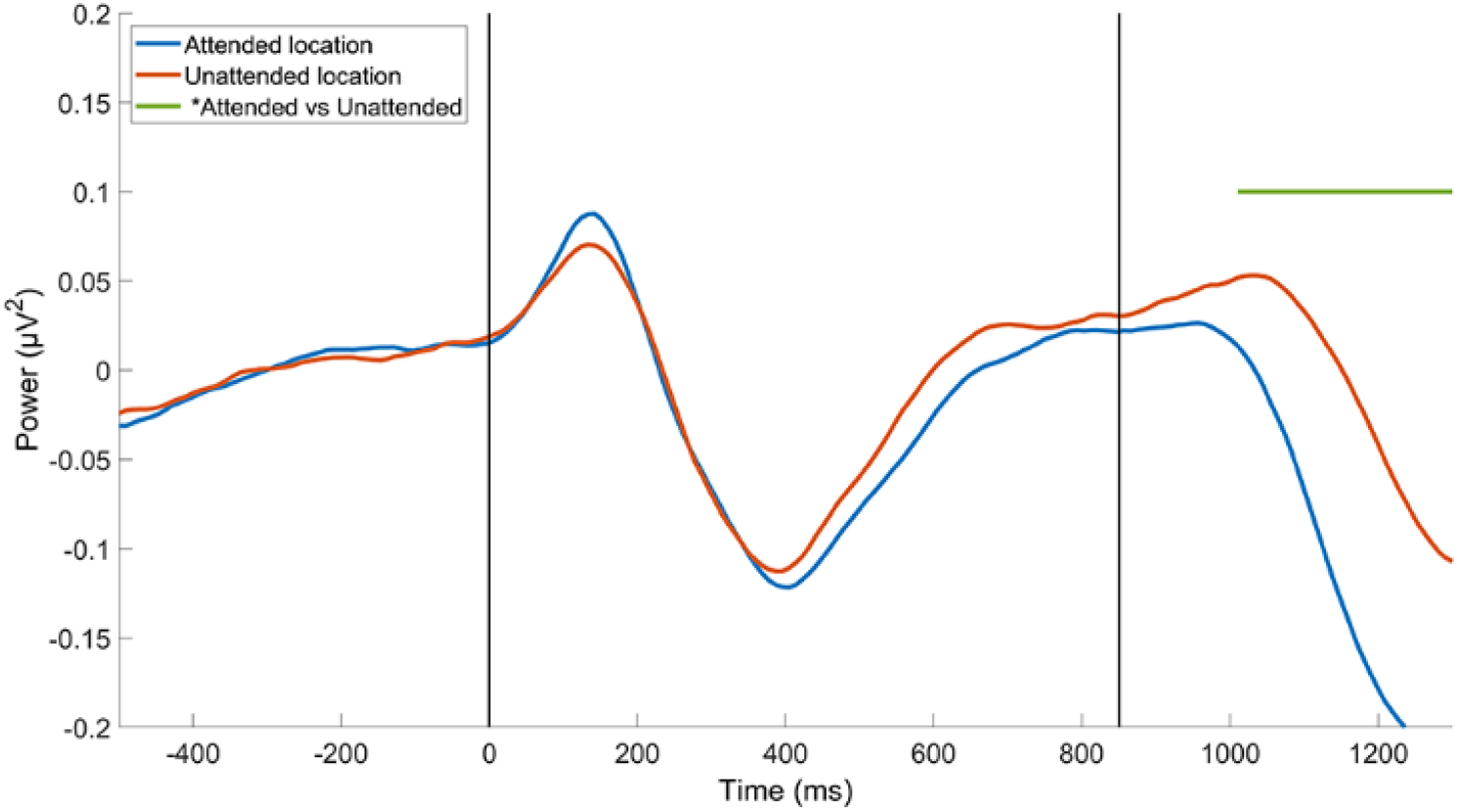
Time course of narrowband alpha power at a priori-defined ROIs, from horizontal blocks. Vertical black lines indicate cue onset (0 ms) and target onset (850 ms). Horizontal line above the data indicates time points with statistically significant comparisons (second-level statistic on cluster-based permutation analysis, p ≤ .01)

#### L1-regularized logistic regression classifier

In order to identify which (and how many) electrodes preferentially represented each of the four target locations (*up, right, down* and *left*) we classified the data at each electrode during the target windows (1000-1200 ms) with a LASSO (Tibshirani, 1996). (The same procedure was followed to create ROIs defined with data corresponding to narrowband alpha power, decomposed alpha power, and 1/f slope). For each of the four target locations, the contribution of each electrode was logistically regressed to classify the tested location against the non-tested locations, using the least square algorithm to select the location-selective electrodes by shrinking the contribution of non-informative or redundant channels to 0 (Tibshirani, 1996).

For participant, each location (the target on ∼100 trials) was classified against the non-target locations (∼100 trials * 3 locations) with a leave-one-trial-out cross-validation procedure. To avoid a bias in the classification for the unbalanced number of samples per class, in each run a down-sampling procedure was applied to each of the three non-tested positions samples to balance the final number of samples per class (∼100 tested location trails vs ∼33 non-tested location trials * 3 positions). The 4 LASSO models were tested on one trial and train on the remaining trials, and repeated until every trial served as a test trial in a classic leave-one-out cross-validation procedure.

In each LASSO logistic regression run, the mean squared error of the logistic regression was estimated for different values of lambda through a 10-fold cross-validation procedure. The classification that showed the minimal cross-validation mean squared error was selected, and electrodes with negative beta coefficients (positive for 1/f slope) were selected for the current tested position in the current leave-one-out cross-validation fold. Consequently, each leave-one-out cross-validation run produced a number of putative sets of electrodes equal to the total number of trials. To verify which electrodes were selected consistently across the cross-validation folds, a bootstrap statistical analysis was implemented. Specifically, electrodes that were selected with a frequency statistically different from 0 across the folds defined the tested location ROI (99% outside the confidence interval). The LASSO procedure on the narrowband alpha power data was able to define one ROI for each of the four tested position in 13 out of 19 subjects, with an average of 7 meaningful electrodes per ROI (up M = 6.79, SD = 5.17; right M = 7.71, SD = 2.93; down M = 7.06, SD = 3.55; left M = 6.88, SD = 3.76; Fig. 2). Correspondingly, the classification procedure on the decomposed alpha power data was successful in 12 out of 19 subjects, with an average of 8 meaningful electrodes per ROIs (up M = 6.19, SD = 3.34; right M = 9.50, SD = 3.61; down M = 7.42, SD = 4.48; left M = 8.47, SD = 3.30; Fig. 3); and the classification procedure on the 1/f slope data was successful in 12 out of 19 subjects, with an average of 10 meaningful electrodes per ROIs (up M = 8.00, SD = 3.92; right M = 11.15, SD = 4.88; down M = 9.89, SD = 4.14; left M = 10.28, SD = 4.20; Fig. 4). For each trial, narrowband alpha power data, the decomposed alpha power datat, and the 1/f slope data from each ROI were relabeled to correspond to its attentional priority state, i.e., *attended, unattended* or *irrelevant* locations. Finally, to simplify the subsequent statistical analysis, activity from the two *irrelevant* locations, i.e., +90 and -90 degrees from the *attended* position, was averaged.

**Figure 2.**
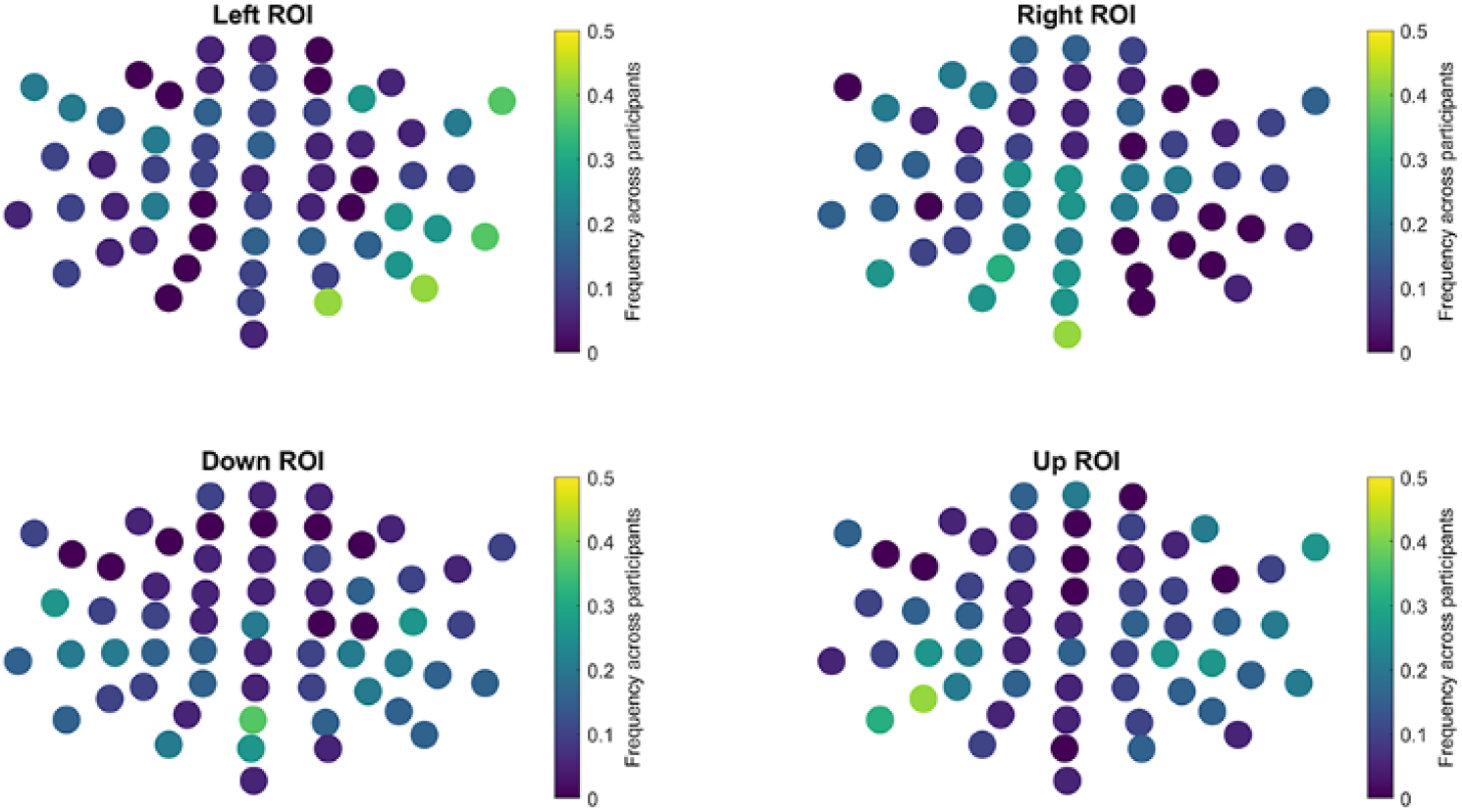
Frequency maps of electrodes selected by multivariate modelling (LASSO on 13/19 participants and mIEM of remaining ones) of narrowband alpha power activity during the target-presentation window. The color represents the frequency which each electrode was selected as a part of that ROI within the sample of subjects.

**Figure 3.**
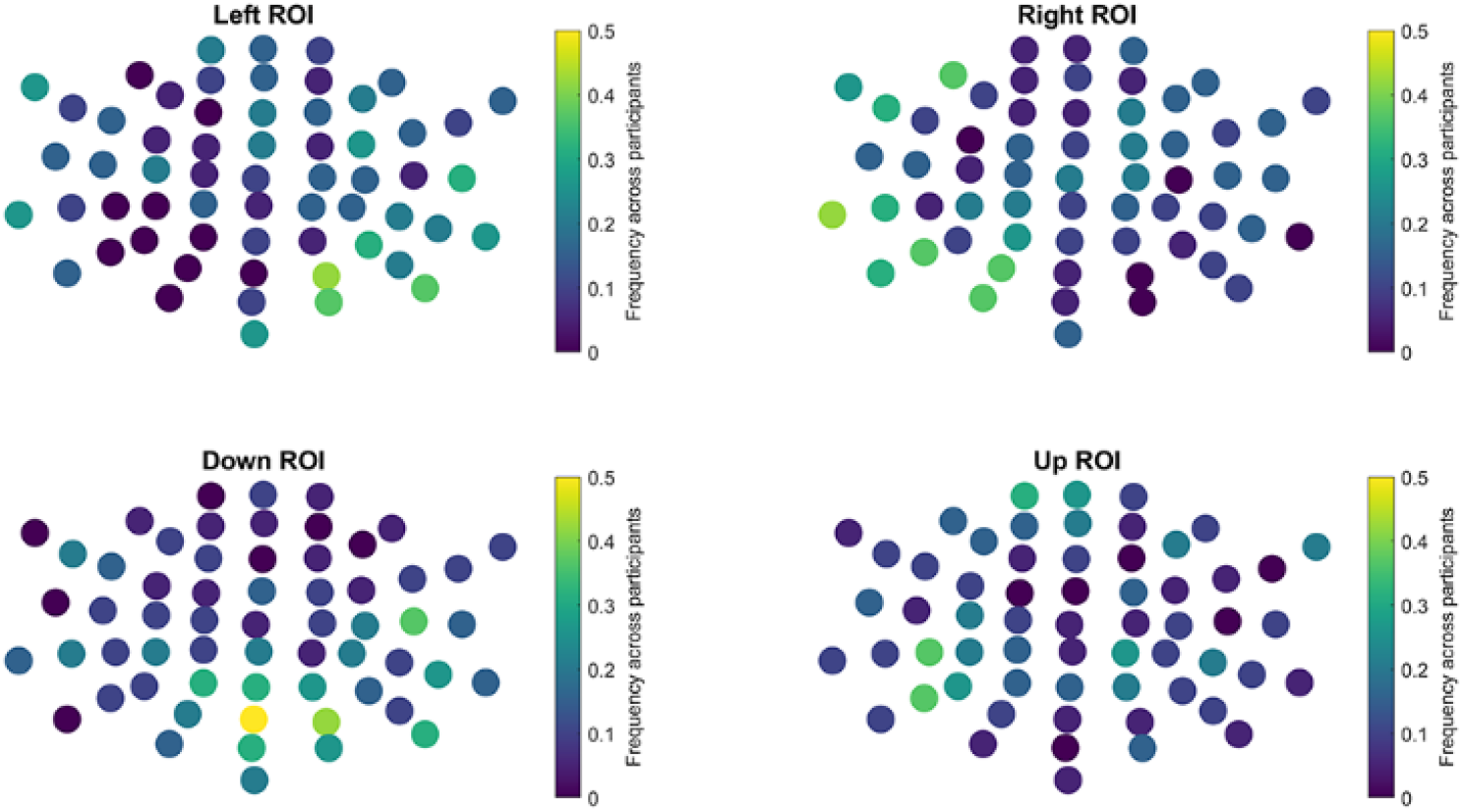
Frequency maps of electrodes selected by multivariate modelling (LASSO on 12/19 participants and mIEM of remaining ones) of the decomposed alpha power component of the EEG (after decomposition) during the target-presentation window. The color represents the frequency which each electrode appeared to be selected as a part of that ROI within the sample of subjects.

**Figure 4.**
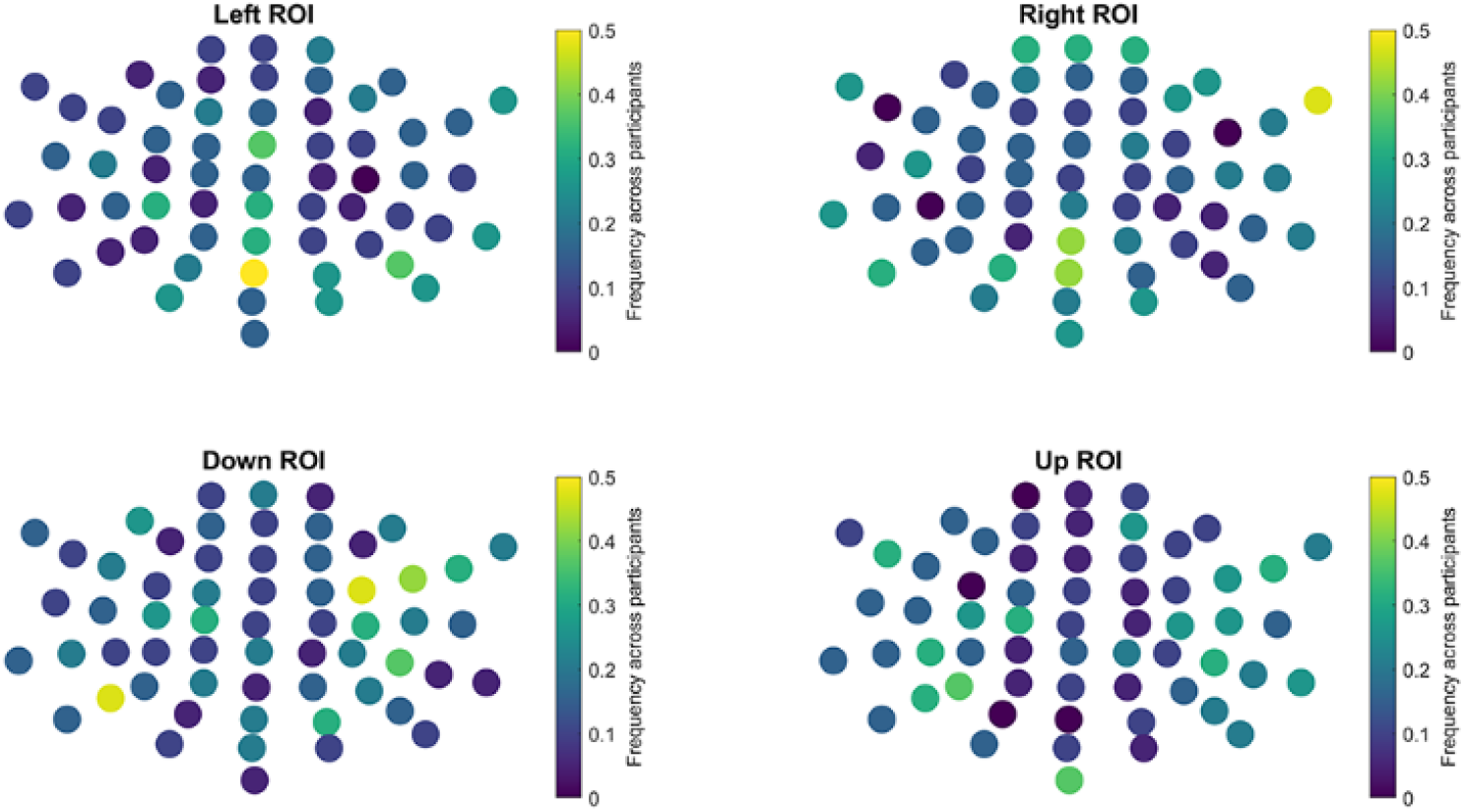
Frequency maps of electrodes selected by multivariate modelling (LASSO on 12/19 participants and mIEM of remaining ones) of 1/f slope component of the EEG (after decomposition) during the target-presentation window. The color represents the frequency which each electrode appeared to be selected as a part of that ROI within the sample of subjects.

#### Multivariate inverted encoding modelling

When the LASSO procedure failed to define all of the four different ROIs, a multivariate Inverted Encoding Modelling (mIEM) was implemented (Sprague and Serences, 2013; Samaha et al., 2016). For each subject, the response in each electrode was modelled as a linear sum of four hypothetical information channels (one per location tested), each corresponding to a delta function centered at its preferred location (i.e., 0°, 90°, 180°, and 270° of polar angle). For all the trials except the one held out for cross validation, a general linear model was solved by regressing the EEG activity from the target window (separately for narrowband alpha power, decomposed alpha power, and 1/f slope) against the basis set of information channels. Next, the resultant matrix containing the mappings of each electrode to the four locations was inverted, and the model tested by feeding it the EEG activity from each of the 60 electrodes from the held-out trial and transforming this into a set of estimated channel responses. This procedure was repeated until every trial served as a test trial in a classical leave-one-out cross-validation procedure. Finally, for each location, we identified the group of electrodes whose activity was most selective for that location. To select the electrodes most selective for each location, beta coefficient matrix values were corrected by subtracting the average across the channels’ beta values. From the average-corrected beta coefficients matrix, the 7, 8 and 10 electrodes with the most negative values (most positive for exponent) were selected for each of the four positions tested, respectively for narrowband alpha power, for decomposed alpha power, and for 1/f slope. The choice of selecting a number of electrodes equal to the average of the ROIs size found in the LASSO procedure is justified by the assumption that this could be the optimal number of electrodes in this experiment for representing one out of four spatial locations.

### Time-frequency series analyses

Evidence for differences between the three attentional priority conditions were assessed statistically using a cluster-based permutation procedure (Maris and Oostenveld, 2007), which was run for each two-condition comparison: *attended* vs *unattended*; *attended* vs *irrelevant*; and *unattended* vs *irrelevant*. Clusters were defined as temporally contiguous timepoints in which two experimental conditions showed a significant difference with a first-level dependent sample *t*-test (p ≤ .05). The second-level statistic was defined as the sum of the *t* statistics of the timepoints in a cluster, obtained by randomly permuting the data between the two experimental conditions within every subject (1000 permutation with Monte Carlo method). Clusters that showed *t* values outside the confidence interval were identified as significant (p ≤ .01).

### Control analysis on stationary spectral activity

Because the fitting-oscillations-and-one-over-f (FOOOF) toolbox (Donoghue et al., 2020b) was originally developed and optimized on frequency series data (Donoghue et al., 2020b), it was important for us to establish the validity of using the method on time-frequency series data, as we intended to do for this experiment. To do this, decomposed alpha power and 1/f slope were derived with FOOOF algorithm also from a Welch’s method spectral distribution (Welch, 1967) as a control analysis. Specifically, a Welch’s method frequency decomposition (1-50 Hz, 0.25 Hz frequencies bins) was run separately on each electrode, each trial, for each subject, across three different time intervals: a baseline interval between -500 and 0 ms, a CTI interval between 350 and 850 ms and a target presentation interval between 850 and 1350 ms. Similarly to the main analysis, FOOOF algorithm (Donoghue et al., 2020b) estimated peak alpha power and exponent values, separately for each electrode, each trial, each subject and each time interval. Finally, the peak alpha power and exponent values were averaged across the electrodes of each ROI and relabeled accordingly to their attentional priority state, i.e., *attended, unattended* and *irrelevant*, and activity from the two *irrelevant* locations, i.e., +90 and -90 degrees from the *attended* position, were averaged.

### Eye-tracking recording and analysis

Eye position across x- and y-axis was recorded monocularly from the right eye with an infrared-based eye tracker with sampling rate of 1000 Hz and a spatial resolution of 0.01° (EyeLink 1000 [SR Research]). Specifically, eye. Movements of the head were limited by a chin rest. Before each experimental block, a standard nine-point-grid calibration was performed, to allow for conversion from raw eye position to gaze position. During the calibration procedure, subjects fixated white dots (1°), serially presented at different locations on the screen, corresponding to a 3×3 array centered on the center of the screen, starting at the upper-left and proceeding in a left-to-right and up-to-down order. Immediately after each calibration procedure, a validation procedure was run, in order to verify the accuracy of the calibration. During validation dots were presented in a random order. X and Y eye gaze data were converted from screen pixels to degree of angle. Blink intervals were identified with a blink detection algorithm and verified by a rater. In order to carry out analyses on micro-saccades, blink intervals were removed from the data, along with samples from 200 ms preceding to 200 ms following each blink. Blink-free gaze position data were then segmented into epochs from -1 to 1999 ms relative to cue onset. Micro-saccades were identified by an expert rater and validated by a secondary rater, blind to the purpose of the study, by visually inspecting the segmented x- and y-gaze data. Eye-movement events larger than 1° of visual angle and outside the CTI were excluded from the analysis. Time, amplitude, and direction of each micro-saccade was recorded. In order to verify that if the direction of the micro-saccades was pointing to the *attended* location, micro-saccades directions were rotated to align the different to-be-attended trial directions on a common *attended* direction, set to 0°.

## Results

### Behavioral performance

Collapsed across axis (horizontal, vertical), mean accuracy was higher for validly than invalidly cued trials (72% correct [SD = 10] vs. 67% [SD = 9], *d* = 0.53), with similar values for horizonal trials (72% correct [11] vs. 66% [12], *d* = 0.52) and vertical trials (71% correct [10] vs. 67% [9], *d* = 0.42). ANOVA with within-subject factors of VALIDITY (validly vs invalidly cued trials) and AXIS (horizontal vs. vertical) confirmed a main effect of VALIDITY (F_(1, 18)_ = 8.85, p < .01), no main effect of AXIS (F_(1,18)_ = 0.00, n.s.), and no interaction (F_(1, 18)_ = 0.59, n.s.). Mean RTs followed a pattern consistent with accuracy: shorter for valid (1004 ms [309]) than invalid (1070 ms [296], *d* = 0.22), with similar values for horizonal trials (1020 ms [331] vs. 1081 ms [294], *d* = 0.19) and vertical trials (988 ms [290] vs. 1059 ms [309], *d* = 0.24). ANOVA revealed a main effect of VALIDITY (F_(1, 18)_ = 4.44, p = .04), no main effect of AXIS (F_(1,18)_ = 1.77, n.s.), and no interaction effect (F_(1, 18)_ = 0.22, n.s.). Visibility ratings followed a pattern consistent with the objective measures, with higher mean ratings for valid (2.27 [0.53] than invalid (2.04 [0.52], *d* = 0.44) trials, and again with similar values for horizonal trials (2.28 [0.53] vs. 2.07 [0.51], *d* = 0.40) and vertical trials (2.27 [0.55] vs. 2.00 [0.54], *d* = 0.50). ANOVA again revealing a main effect of VALIDITY (F_(1, 18)_ = 6.59, p = .02), no effect of AXIS (F_(1,18)_ = 1.53, n.s.), and no interaction (F_(1, 18)_ = 1.93, n.s.).

The mean percentage of trials in which one or more microsaccades were detected was 46% (right cued (45% [20]); down cued (43% [16]); left cued (47% [18]); up cued (48% [18]); values that did not differ (*F*_(3, 45)_ = 2.05, n.s.)), and they occurred an average of 405 ms after the cue presentation. A second microsaccade was detected on 41% of these trials (right cued (39% [22]); down cued (41% [19]); left cued (43% [21]); up cued (39% [22]); values that did not differ (*F*_(3, 45)_ = 0.55, n.s.)), they occurred an average of 573 ms after cue presentation, and 63% of them were a return to fixation. In order to test if micro-saccades executed during the CTI were biased toward the attended location, violation of a uniform distribution of micro-saccades around the attended location (0 DVA) was verified with a V-test (Zar, 1999), run with the CircStat toolbox (Berens, 2009) on Matlab (2020b, The MathWorks, Natick, MA). The distribution of the direction of micro-saccades during the CTI was significantly nonuniform, and biased toward the attended position (M = +5.62°, AD = 0.94°; V_(1, 18)_ = 6.60, p = 0.01, *d* = 5.98). To summarize, micro-saccades were biased toward the *attended* location from ∼400 ms after the spatial cue presentation, suggesting the influence of spatial expectations also on the control of eye-movements.

### Narrowband alpha power

Narrowband alpha power recorded from the electrodes representing the *attended* location was significantly lower compared to electrodes representing the *unattended* location, starting from 386 ms after cue presentation and lasting until the end of the trial (p > .001), and it differed from the electrodes representing the *irrelevant* locations starting from 84 ms before cue presentation and lasting until the end of the trial (p > .001). Of primary importance for the question that we set out to address, narrowband alpha power did not differ between electrodes representing the *unattended* and the *irrelevant* locations at any point during the trial (Fig. 5).

**Figure 5.**
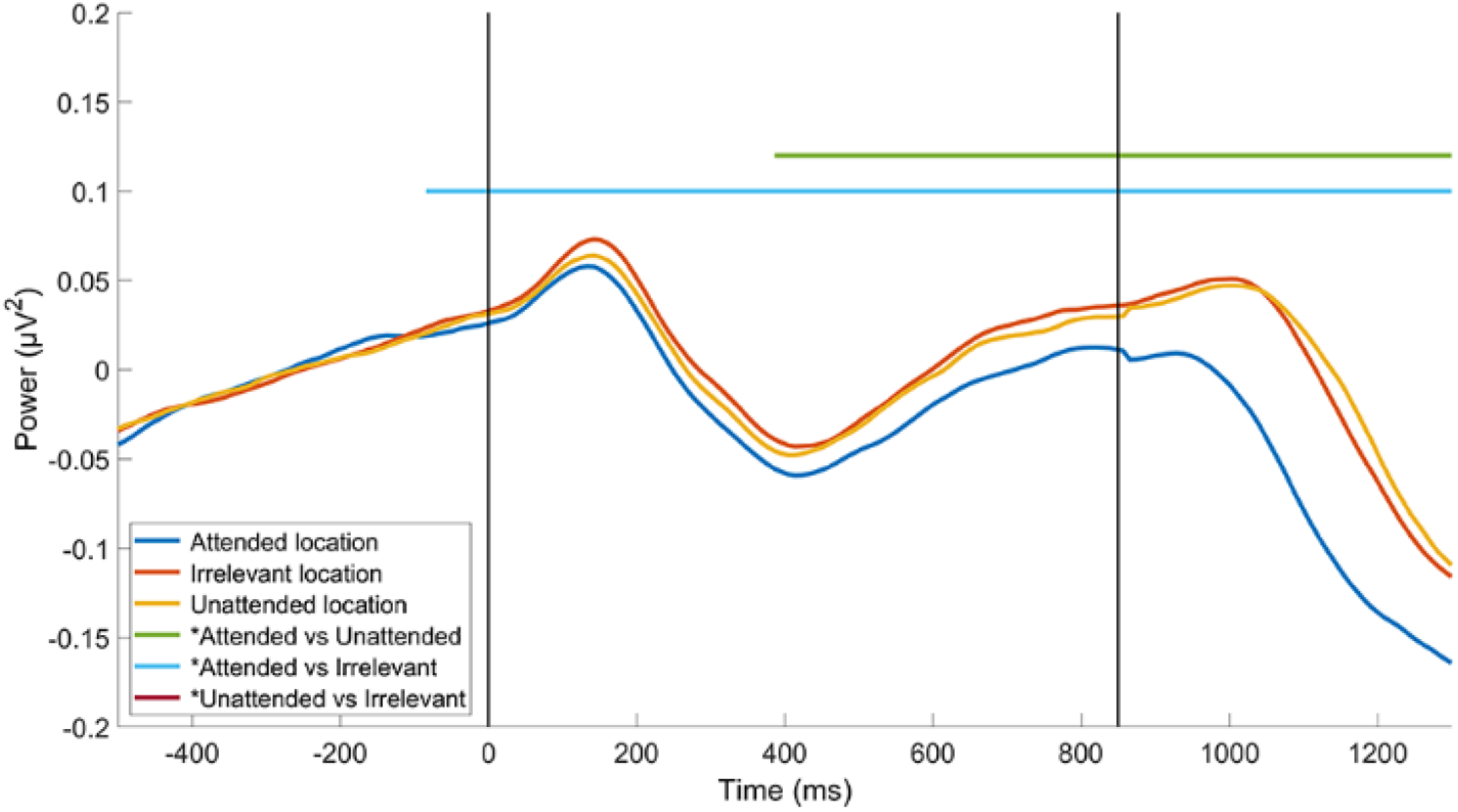
Time course of narrowband alpha power across ROIs derived from multivariate analyses of narrowband alpha-power data (see Fig. 2). Vertical black lines indicate cue onset (0 ms) and target onset (850 ms). Horizontal lines above the data indicate time points with statistically significant comparisons (second-level statistic on cluster-based permutation analysis, p ≤ .01)

Following the logic of our design, the absence of evidence for any difference between ROIs representing *unattended* and *irrelevant* locations is consistent with secondary-consequence accounts of attention-related alpha dynamics: When attention selects a location, the physiological state of all other locations returns to a common baseline level. Nonetheless, the theoretical implications of an absence of evidence between two conditions can be difficult to interpret unless one can rule out the less interesting possibility that the experiment simply lacks the sensitivity to detect a difference. Thus, the interpretability of these results with narrowband alpha would be clearer if a difference between the *unattended* and *irrelevant* ROIs could be found with some other physiological signal – or with a component of the original signal. This provides motivation for the decomposition of these same EEG data into periodic and aperiodic components, to determine whether either of these components might discriminate the *unattended* from the *irrelevant* location.

### Decomposed EEG signals

The idea motivating the decomposition of the EEG into periodic and aperiodic components is that the conventional method of bandpass filtering leaves uncertain whether attention-related effects like those illustrated in Fig. 5 are due to effects of attention on true oscillations in the 8-14 Hz range, on aperiodic components that also influence this range, or on some mixture of the two. To address these questions comprehensively, we examined the dynamics of the decomposed alpha and 1/f slope signals in each of two sets of ROIs: the ROIs generated with narrowband alpha data (Fig. 2) and the ROIs generated with the decomposed alpha and 1/f slope signals, respectively (Fig. 3 and 4). The reason for doing so in the narrowband-alpha ROIs is to assess the extent to which the pattern observed in Fig. 5, (which broadly replicates a well-established finding in the literature for attended vs. unattended electrodes), is due to “true” alpha-band oscillations or to aperiodic elements in the narrowband signal. Additionally, however, it is important to note that the narrowband-alpha ROIs may not be selective for up, right, down, and left for either the decomposed alpha or the 1/f slope signals. Stated another way, the topography of retinotopy for either of these two putatively independent generators may be different that it is for the scalp-level EEG signal that is a mixture of the two. Thus, a more valid way to assess the effects of spatial cuing on decomposed alpha is to identify electrodes that are selective for the representation of these four locations by decomposed alpha, and the same is true for 1/f slope.

#### Decomposed alpha power at ROIs spatially selective for narrowband alpha

In the ROIs identified by regressing target related narrowband alpha power, decomposed alpha power discriminated between *attended* and both *unattended* and *irrelevant* locations, but only during target presentation (Fig. 6). Specifically, decomposed alpha power recorded from the electrodes representing the *attended* location was significantly lower compared to electrodes representing the *unattended* location, starting from 929 ms after cue presentation and lasting until the end of the trial (p < .001). Furthermore, the decomposed alpha power recorded from the *attended* location was also significantly lower compared to electrodes representing the task-*irrelevant*, starting from 812 ms and lasting until the end of the trial (p < .001). Finally, no significant difference was found between *unattended* and *irrelevant* locations across all the trial duration (all p > .01). Overall, decomposed alpha power at ROIs spatially selective for narrowband alpha failed to show a modulation of any of the ROIs during the CTI, suggesting a lack of correspondence between narrowband and decomposed alpha power distribution across the scalp.

**Figure 6.**
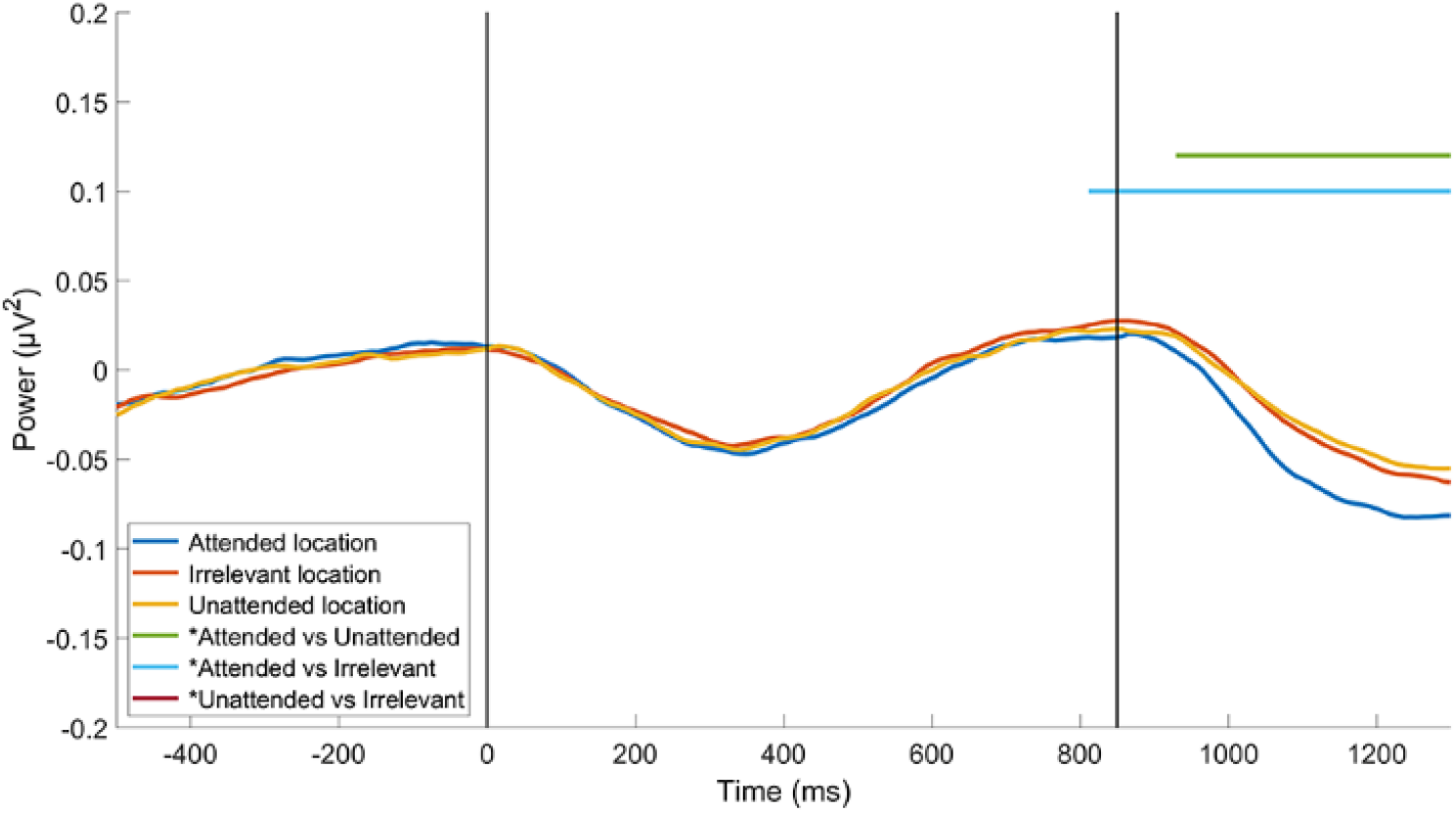
Time course of decomposed alpha power across ROIs derived from multivariate analyses of narrowband alpha power. Vertical black lines indicate cue onset (0 ms) and target onset (850 ms). Horizontal lines above the data indicate time points with statistically significant comparisons (second-level statistic on cluster-based permutation analysis, p ≤ .01)

#### Decomposed alpha power at ROIs spatially selective for decomposed alpha power

In the signal-selective ROI, decomposed alpha power only discriminated between the three attentional priority states during target presentation (Fig. 7). As was the case with narrowband alpha, the divergence of decomposed alpha power corresponding to the *attended* versus the *irrelevant* location began earlier (694 ms after cue presentation; p < .001) than the divergence of *attended* to unattended (902 ms after cue presentation; p < .001). Unlike narrowband alpha, however, decomposed alpha power corresponding to the *unattended* location did eventually diverge from decomposed alpha power corresponding to the *irrelevant* location, taking on a higher value starting from 1083 ms after cue presentation (p < .001). Note that because all of these effects occurred shortly before, or after, the onset of the target, it is unlikely that they reflect an important contribution to a cue-triggered anticipatory shift of spatial attention.

**Figure 7.**
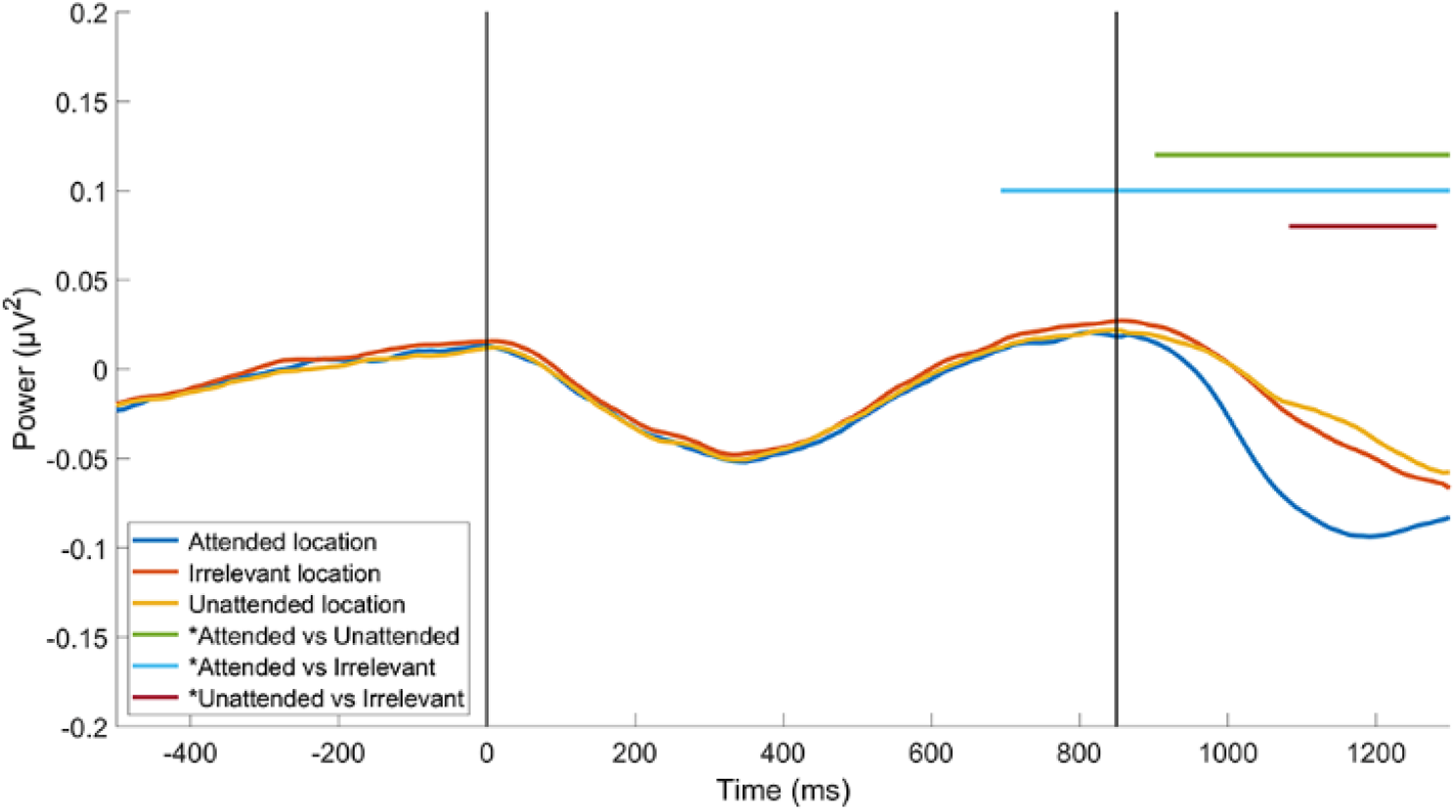
Time course of decomposed alpha power across ROIs derived from multivariate analyses of decomposed alpha power. Vertical black lines indicate cue onset (0 ms) and target onset (850 ms). Horizontal lines above the data indicate time points with statistically significant comparisons (second-level statistic on cluster-based permutation analysis, p ≤ .01)

#### 1/f slope at ROIs spatially selective for narrowband alpha

In the ROIs identified by regressing target related narrowband alpha power, 1/f slope discriminated between *attended* and both *unattended* and *irrelevant* locations, but only during target presentation (Fig. 8). Specifically, 1/f slope from the *attended* location was significantly lower compared to electrodes representing the *unattended* location, starting from 1002 ms to 1191 ms (p < .001), and similarly was significantly lower compared to *irrelevant* location, starting from 1002 ms to 1173 ms (p < .001). Finally, no significant difference was found between *unattended* and *irrelevant* locations across all the trial duration (all p > .01). In the same way as for the decomposed alpha power, 1/f slope at ROIs spatially selective for narrowband alpha failed to show a modulation of any of the ROIs during the CTI, suggesting a lack of correspondence between narrowband and 1/f slope distribution across the scalp.

**Figure 8.**
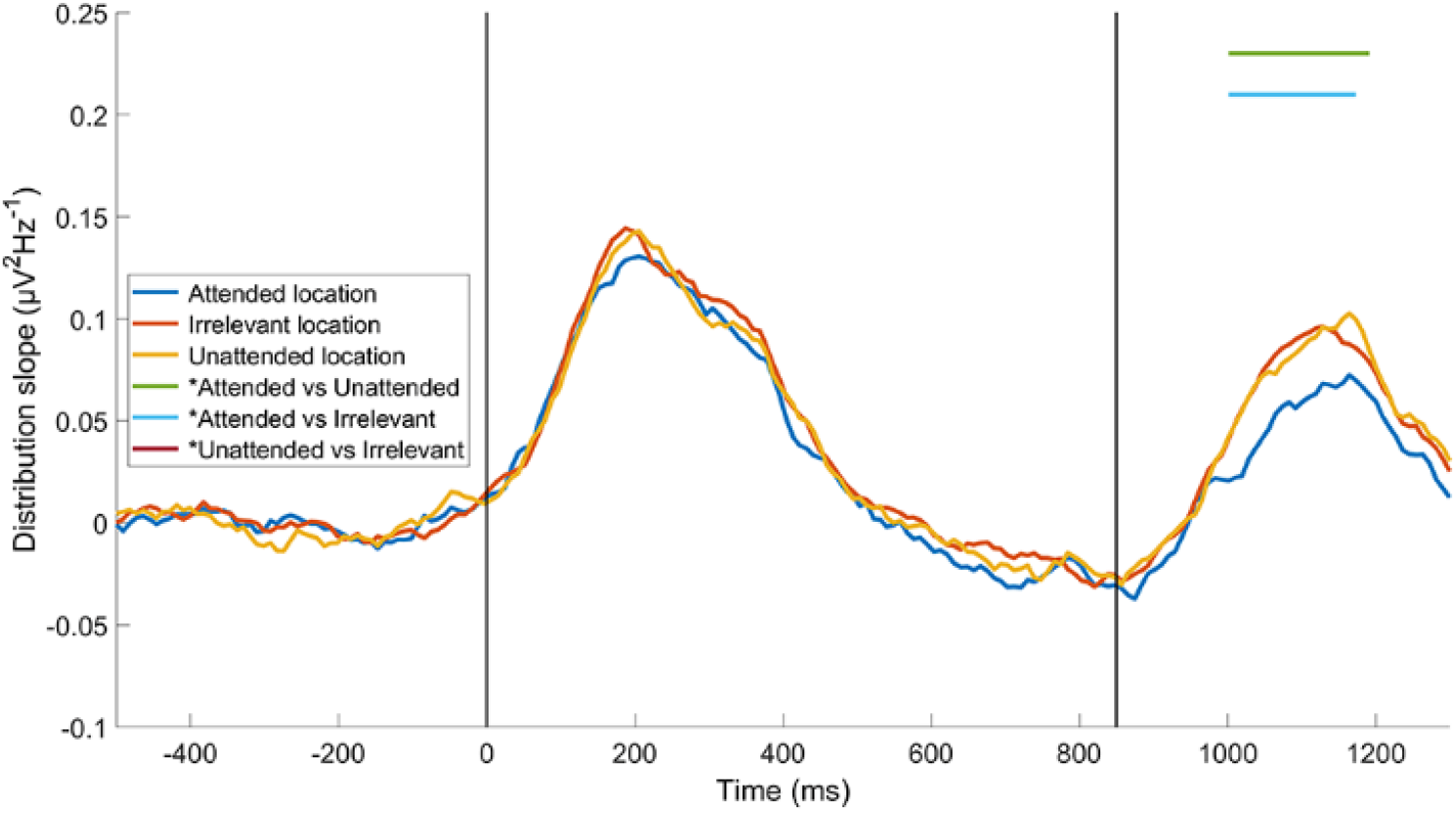
Time course of 1/f slope across ROIs derived from multivariate analyses of narrowband alpha power. Vertical black lines indicate cue onset (0 ms) and target onset (850 ms). Horizontal lines above the data indicate time points with statistically significant comparisons (second-level statistic on cluster-based permutation analysis, p ≤ .01)

#### 1/f slope at ROIs spatially selective for 1/f slope

In a pattern markedly different from narrowband alpha and decomposed alpha, 1/f slope differed at the *irrelevant* versus *unattended* ROI during the pretrial ITI and for the entirety of the trial (p < .001), suggesting that 1/f slope was tonically elevated at *irrelevant* ROIs for the entirety of the block. (Recall that, because the factor AXIS was blocked, electrodes corresponding to an *irrelevant* location retained this status for the duration of each block.) 1/f slope also differed between *irrelevant* and *attended* ROIs during the pretrial ITI and for the entirety of the trial (p < .001). 1/f slope did not differ between *attended* and *unattended* ROIs until a window beginning after target onset (from 884-1255 ms after cue presentation; p < .001), during which 1/f slope was greater at the *unattended* ROI (Fig. 9). From a physiological perspective, this result suggests that locations that are potential targets of selection (i.e., *attended* or *unattended*) may differ from locations that are known to be task-*irrelevant* in that the E:I in the former is tonically elevated. From an interpretational perspective, this dissociation of *irrelevant* from *unattended* suggests that the null findings with the narrowband alpha and decomposed alpha signals can, indeed, be interpreted as evidence that attention-related increases in alpha-band power do not reflect an active-suppression mechanism, but may instead simply be a consequence of the withdrawal of attention from the unattended region.

**Figure 9.**
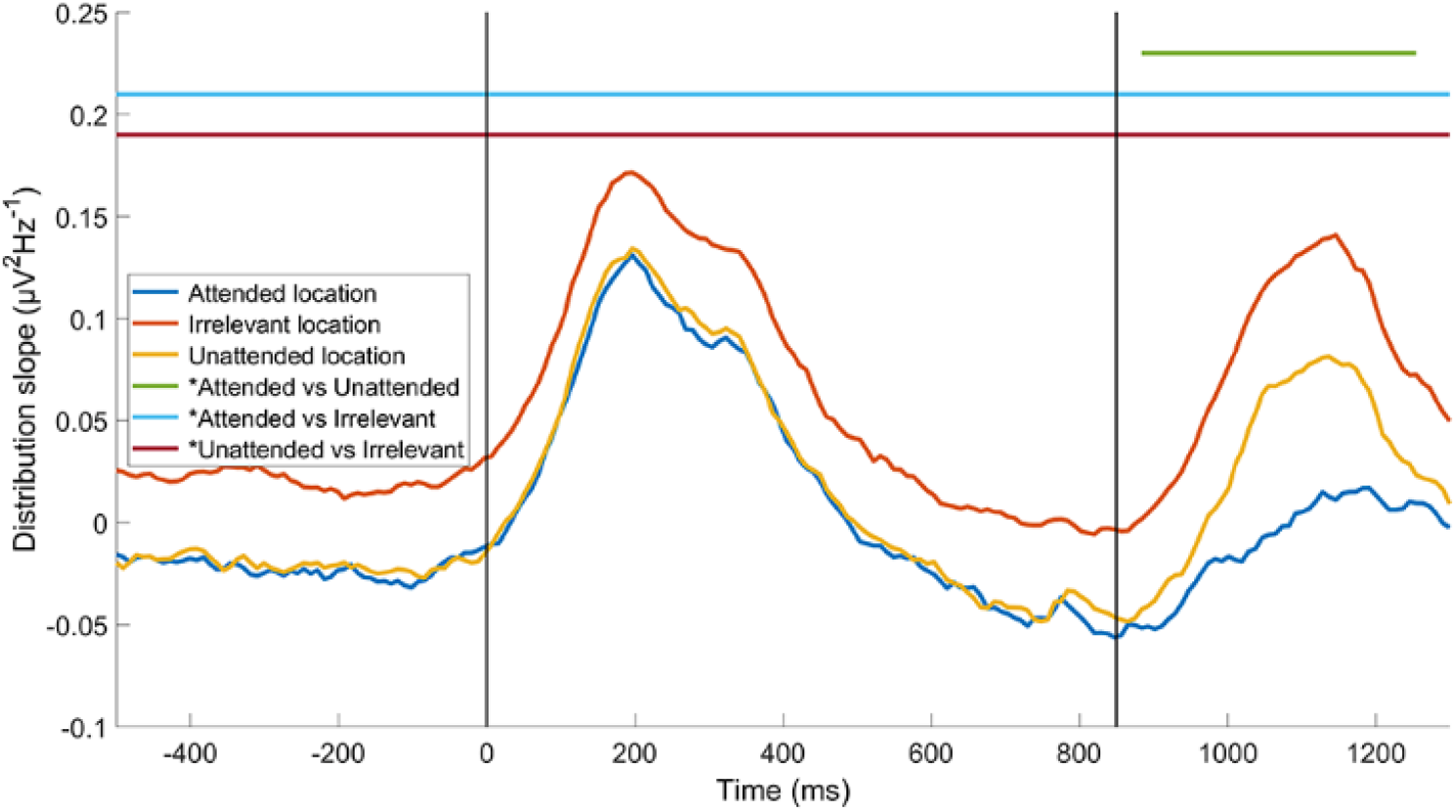
Time course of 1/f slope across ROIs derived from multivariate analyses of the 1/f slope data. Vertical black lines indicate cue onset (0 ms) and target onset (850 ms). Horizontal lines above the data indicate time points with statistically significant comparisons (second-level statistic on cluster-based permutation analysis, p ≤ .01)

### Control frequency decomposition

Possible activity difference between the three attentional priority conditions across the three time intervals were statistically assessed running repeated-measures ANoVAs. Specifically, two separate ANoVAs for decomposed alpha power and 1/f slope were run. Each ANoVA had TIME (baseline vs CTI vs target presentation time intervals) and PRIORITY (*attended* vs *unattended* vs *irrelevant* locations) as within-subject factors. All post-hoc comparisons were conducted using the Newman– Keuls test. These statistical analyses were performed using STATISTICA (Version 12.0; StatSoft).

Regarding the ANoVA run for the decomposed alpha power (Fig. 10), a significant TIMExPRIORITY interaction effect (F_4,72_ = 52.30, p < .01) was found. Specifically, for the CTI time interval, the *attended* ROI (M = 6.16 µV^2^ [0.24]) showed a significant lower decomposed alpha power compared to both the *irrelevant* (M = 6.20 µV^2^ [0.24], p = .04, *d* = 0.17) and *unattended* (M = 6.19 µV^2^ [0.24], p = .04, *d* = 0.13) ROIs, whereas no significant difference was found between *irrelevant* and *unattended* ROIs (p = .82). Moreover, for the target presentation time interval, the *attended* ROI (M = 5.83 µV^2^ [0.19]) showed a significant lower decomposed alpha power compared to both the *irrelevant* (M = 6.02 µV^2^ [0.20], p < .01, *d* = 0.97) and *unattended* (M = 6.05 µV^2^ [0.20], p < .01, *d* = 1.13) ROIs, and the *irrelevant* ROI showed a significant lower decomposed alpha power compared to the *unattended* (p = .02, *d* = 0.15) ROI. Finally, no significant post-hoc tests were found significant for the baseline time interval (all ps > .11). Additionally, significant main effects of both TIME (F_2,36_ = 4.97, p = .01) and PRIORITY (F_2,36_ = 40.7, p < .01) were found, described by a significant lower decomposed alpha power during target presentation (M = 5.96 µV^2^ [0.20]) compared to both baseline (M = 6.15 µV^2^ [0.25], p = 0.02, *d* = 0.84) and CTI (M = 6.18 µV^2^ [0.24], p = .02, *d* = 0.96) time intervals, and by a lower decomposed alpha power in the *attended* ROI (M = 6.05 µV^2^ [0.23]) compared to both *irrelevant* (M = 6.12 µV^2^ [0.23], p < .01, *d* = 0.30) and *unattended* ROIs (M = 6.13 µV^2^ [0.23], p < .01, *d* = 0.35). Overall, decomposed alpha power, estimated on frequency series, showed a similar modulation of ROIs representing *unattended* and *irrelevant* locations, suggesting that spatial expectations modulations on true periodic alpha power don’t discriminate between *unattended* and *irrelevant* locations.

**Figure 10.**
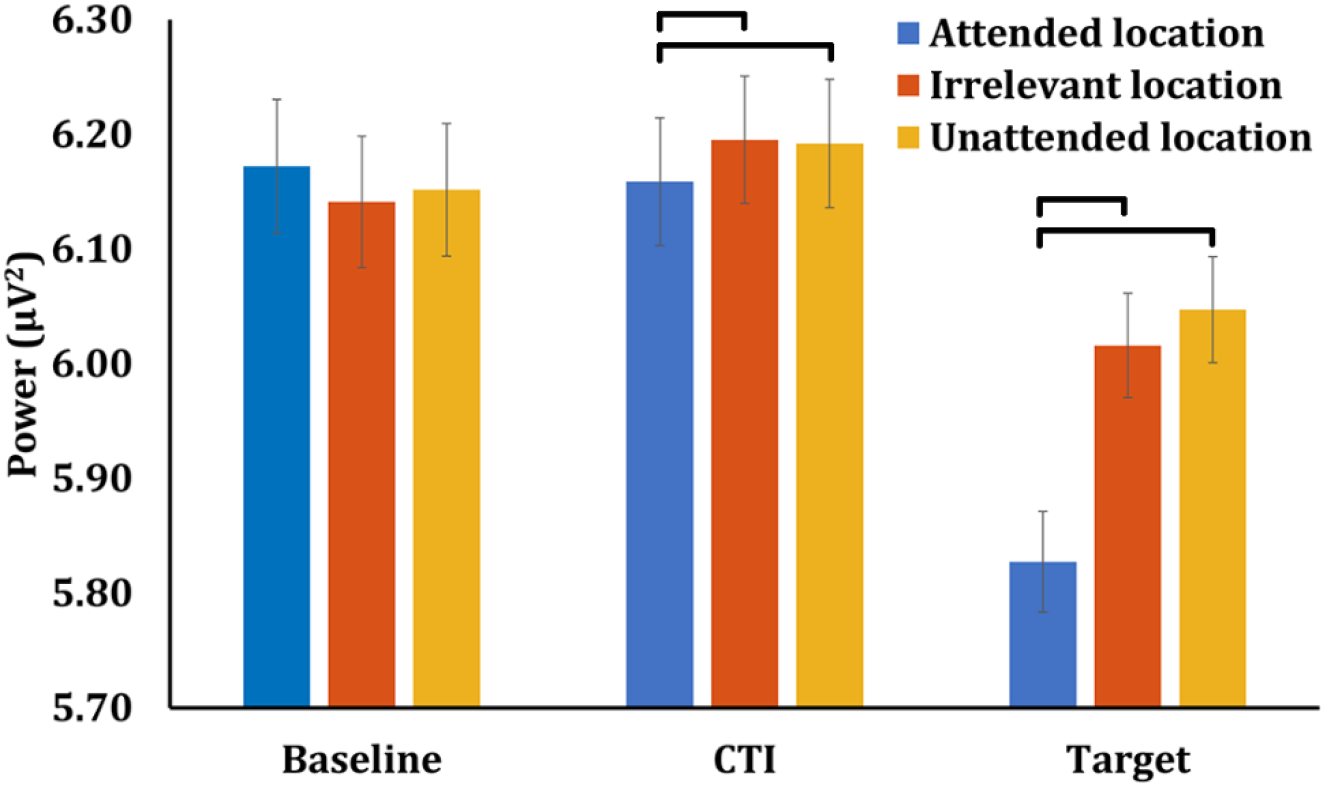
Decomposed alpha power across baseline (−500-0 ms), CTI (350-850 ms), and target presentation (850-1350 ms) epochs, from ROIs derived from multivariate analyses of decomposed alpha power. Black lines indicate statistically significant comparisons (Newman–Keuls, p ≤ .05)

Regarding the ANoVA run for the 1/f slope (Fig. 11), a significant TIMExPRIORITY interaction effect (F_4,72_ = 23.67, p < .01) was found. Specifically, for the baseline time interval, the *irrelevant* ROI (M = 5.84 µV^2^Hz^-1^ [0.20]) showed a significant higher 1/f slope compared to both the *attended* (M = 5.69 µV^2^Hz^-1^ [0.20], p < .01, *d* = 0.75) and *unattended* (M = 5.71 µV^2^Hz^-1^ [0.20], p < .01, *d* = 0.65) ROIs, whereas no significant difference was found between *attended* and *unattended* ROIs (p = .37). For the CTI time interval, similarly to the baseline time interval, the *irrelevant* ROI (M = 5.86 µV^2^Hz^-1^ [0.20]) showed a significant higher 1/f slope compared to both the *attended* (M = 5.75 µV^2^Hz^-1^ [0.20], p < .01, *d* = 0.55) and *unattended* (M = 5.74 µV^2^Hz^-1^ [0.20], p < .01, *d* = 0.60) ROIs, whereas no significant difference was found between *attended* and *unattended* ROIs (p = .89). Moreover, for the target presentation time interval, the *irrelevant* ROI (M = 6.18 µV^2^Hz^-1^ [0.20]) showed a significant higher 1/f slope compared to both the *attended* (M = 5.90 µV^2^Hz^-1^ [0.20], p < .01, *d* = 1.4) and *unattended* (M = 6.00 µV^2^Hz^-1^ [0.20], p < .01, *d* = 0.90) ROIs, and the *unattended* ROI showed a significant higher 1/f slope compared to the *attended* (p = .02, *d* = 0.5) ROI. Additionally, significant main effects of both TIME (F_2,36_ = 33.7, p > .01) and PRIORITY (F_2,36_ = 45.9, p < .01) were found, described by a significant higher 1/f slope during target presentation (M = 6.02 µV^2^Hz^-1^ [0.20]) compared to both baseline (M = 5.75 µV^2^Hz^-1^ [0.20], p < .01, *d* = 1.35) and CTI (M = 5.78 µV^2^Hz^-1^ [0.20], p < .01, *d* = 1.20) time intervals, and by a higher 1/f slope in the *irrelevant* ROI (M = 5.96 µV^2^Hz^-1^ [0.20]) compared to both *attended* (M = 5.78 µV^2^Hz^-1^ [0.20], p < .01, *d* = 0.9) and *unattended* ROIs (M = 5.82 µV^2^Hz^-1^ [0.20], p < .01, *d* = 0.70). Overall, 1/f slope, estimated on the frequency series, was tonically higher in ROI representing the *irrelevant* locations compared to both *attended* and *unattended* locations across the duration of both the trial and the ITI time interval, corroborating that expectations about the spatial configuration of the current block modulate the aperiodic component of the spectral distribution tonically.

**Figure 11.**
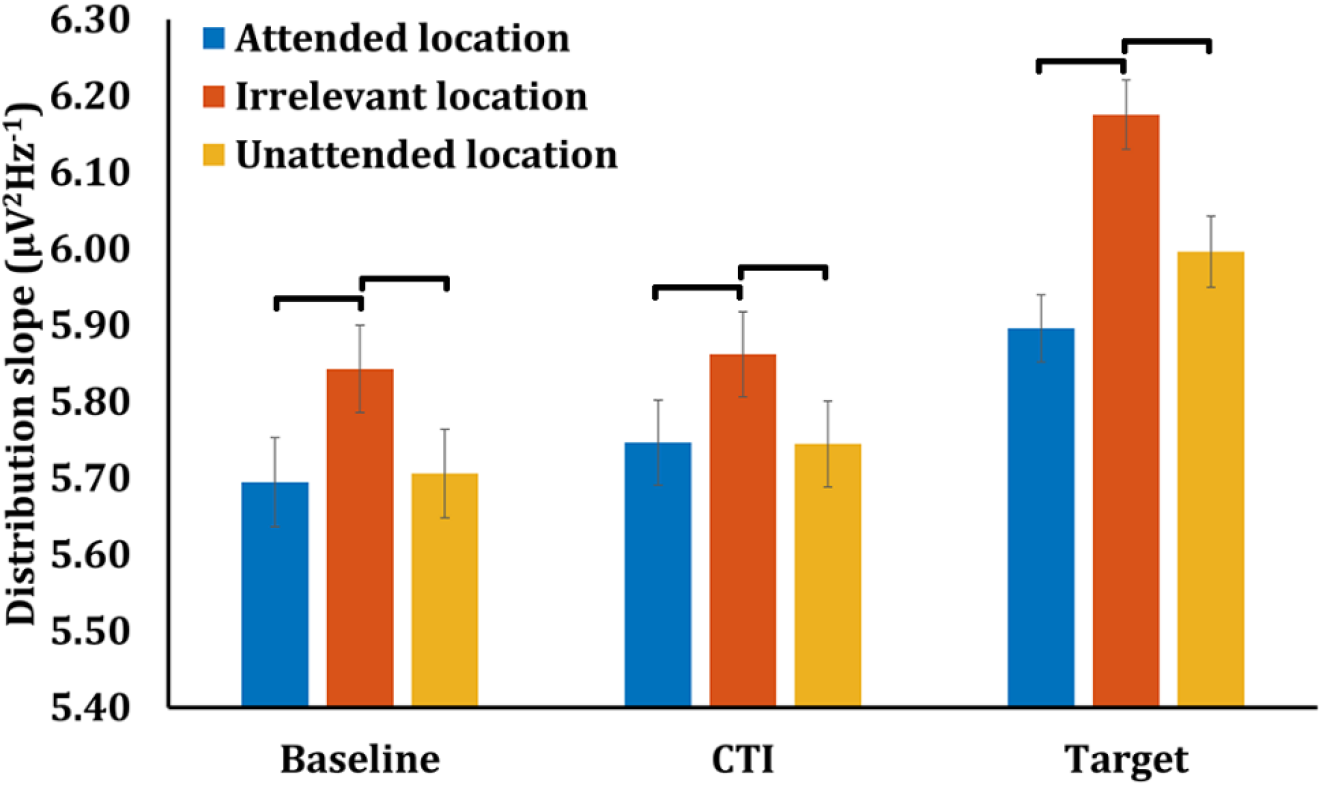
1/f slope across baseline (−500-0 ms), CTI (350-850 ms,) and target presentation (850-1350 ms) ROIs derived from multivariate analyses of 1/f slope data. Black lines represent statistically significant comparisons (Newman–Keuls, p ≤ .05).

## Discussion

In this experiment we blocked trials such that, in any given block, targets would only appear in two of four possible locations, thereby rendering the other two locations *irrelevant* on that block. Behaviorally, the spatial information conveyed by the symbolic cue influenced performance in ways that are well established in the literature: improved discrimination on validly cued trials, and a biased pattern of microsaccades during the CTI. Starting with the narrowband alpha signal (i.e., the signal comparable to that used in the vast majority of studies of spatial attention to date), our results replicated the canonical pattern of the cue-triggered dissociation of alpha power at *attended* versus *unattended* locations. Of principal theoretical interest, they also showed that the dynamics of alpha power at *irrelevant* locations very closely matched those of *unattended* locations. This finding is difficult to reconcile with interpretations of alpha as a mechanism for the implementation of attention control (e.g., Kelly et al., 2006; Jensen and Mazaheri, 2010; Sadaghiani and Kleinschmidt, 2016), and instead offers support for the possibility that attention-related changes in alpha power may be consequences of some other factors (Antonov et al., 2020). For example, if selection of the attended location is accomplished via increased input (via spiking) from source areas, this increased input may cause a decrease in alpha power; concomitantly, a decrease in top-down input at non-selected areas, combined with lateral inhibition from the selected location, would result in an increase. Although our empirical result can be summarized as a failure to reject the null hypothesis (that *unattended* and *irrelevant* do not differ), the success at dissociating *unattended* from *irrelevant* locations with an aperiodic (i.e., non-oscillatory) component of the same signal suggests that the null finding with narrowband alpha is not due to a general lack of sensitivity with this design. Furthermore, the facts that *attended* and *unattended* did not differ with regard to 1/f slope, and that this effect persisted throughout the block, including during the ITI, suggests that physiological state (perhaps E:I balance) is modified at locations where selection is predicted to occur.

A convention for EEG studies of Posner-style cuing tasks is to cue targets to the right or left of fixation, then label signals from a predefined set of electrodes contralateral to the cue as “*attended*” and from a predefined set of electrodes ipsilateral to the cue as “*unattended*.” One limitation of this approach is that it doesn’t allow for the distinction between “*unattended*” and “*irrelevant*.” This is important because if the increase in alpha power at *unattended* electrodes is truly a mechanism deployed to inhibit processing at the *unattended* location, one should expect this signal to be retinotopically focused. Indeed, for studies that cue multiple locations equally spaced around fixation, the scalp distribution of attention-related increases in alpha-band power is relatively focused, certainly not at every electrode except those corresponding to the cued location (c.f., Samaha et al., 2016). To be able to create electrodes selective for *irrelevant* locations, it was necessary for us to use an analytic method for identifying electrodes selective for each of the four target locations. The validity of this approach is demonstrated by the fact that it generates results with narrowband alpha (Fig. 5) that are qualitatively similar to those generated with the conventional lateralized procedure (Fig. 1). (Note that quantitative comparison of figures 1 versus 5 is complicated by the fact that Fig. 5 was generated with twice as many trials (because it also includes data from *up*/*down* blocks).)

These results are consistent with the idea that spatial selection is achieved, at least in part, via an increase in action potentials originating from cortical circuits in the dorsal attention network (DAN; e.g., Corbetta and Shulman, 2002) and the pulvinar nucleus (e.g., Saalmann and Kastner, 2011). Although an effect of this activity is to synchronize local field potentials of targeted circuits with those generating these afferent signals (e.g., Levichkina et al., 2021), the resulting spike-field coherence, whether prominent in the alpha band (e.g., Saalmann et al., 2012) or in other frequency bands (e.g., Mendoza-Halliday et al., 2014) are likely too small to detect with scalp EEG. Our results also suggest that the cue-locked macro-scale increase in alpha-band power that is observed at *unattended* locations may be a secondary consequence of selection happening elsewhere, a return to an “idling” state similar to what is observed upon eye closure. We note that this interpretation is consistent with recent work that has concluded from careful measurement of the timing of attention-related changes in steady-state evoked potentials that changes in alpha-band power measured at the scalp are too slow for them to have a direct role in attentional control (Antonov et al., 2020).

Importantly, our results are in line with the longstanding idea that spatial expectation about a forthcoming visual stimulus modulates alpha power in visual areas representing the attended location (e.g., Thut et al., 2006; Rihs et al., 2009). There is broad consensus that this decrease in alpha power does not correspond to merely a residual effect of the visual processing of the cue, but that it reflects an endogenous modulation of posterior alpha that has the effect of increasing the excitability of visual circuits corresponding to the cued location. Furthermore, our results with decomposed components of the EEG signal confirm that this effect at the target of selection is truly specific to oscillations in the alpha band, because *attended-unattended* differences were observed in only the decomposed alpha signal, not in the aperiodic component of the EEG, which is also prominent at 8-14 Hz.

One novel aspect of our results is that they include a heretofore undescribed component of the EEG signal that does differentiate *irrelevant* from *unattended* locations: the slope of the aperiodic 1/f-like component. Because this measure is believed to reflect local cortical E:I balance (Freeman and Zhai, 2008; Voytek et al., 2015; Gao et al., 2017; Donoghue et al., 2020b; Ostlund et al., 2021), this aspect of our findings gives us license to propose an attention-related mechanism that is different from shifts of selective attention. Knowledge that behaviorally relevant targets will only occur at specific locations in the visual field may prompt a tonic change in E:I balance at these locations, preconditioning them for anticipated target processing.

We conclude with some methodological considerations. The combined use of univariate and multivariate analysis techniques allowed us to capitalize on the respective advantages of each, while avoiding many of their respective shortcomings. Univariate methods can produce measures that relate directly to brain physiology, such as functional differences between desynchronization and synchronization of oscillations in the alpha band (Thut et al., 2006; Romei et al., 2008b, 2008a; Rihs et al., 2009; Capilla et al., 2014). A shortcoming of this approach, however, is that they often require making a priori assumptions about “how the brain works,” as one does when selecting electrodes for a priori-defined ROIs. If any of these assumptions are incorrect, interpretation of the data will necessarily be flawed. Conversely, although multivariate techniques offer improved specificity, for example by weighting the contribution of various kinds of information across different features, they often produce measures without clear correspondence to observable physiological mechanisms. In the current study we used multivariate methods (Lasso and mIEM) to select electrodes whose activity was selective for only one of the four locations. This optimized the specificity of the signals that we measured and didn’t require us to assume that we know how *up, right, down*, and *left* are represented at the scalp. In particular, the loss function of the Lasso regularization procedure maximizes the contribution of location-informative channels while pushing to zero the contribution of non-relevant and redundant channels (Tibshirani, 1996). Having so identified the ROIs, we then used a univariate approach (i.e., averaging across multiple electrodes to extract one signal per ROI) to assess fluctuations in physiological states (e.g., synchronization of LFPs, E:I balance) that are directly measurable. The improved sensitivity of this approach is evidenced by the presence of an earlier modulation of alpha power in the attended position in comparison to what has been observed using conventional lateralization procedures (Thut et al., 2006; Rihs et al., 2009). In our analyses, signals from a priori-defined lateralized ROIs did not show a statistically reliable cue-related modulation of alpha power until 1000 ms post cue, whereas from multivariate analysis-derived ROIs the same modulation was evident 380 ms after cue onset. The results with these multivariate analysis-derived ROIs are decidedly better aligned with behavioral estimates of the latency of endogenously triggered shifts of attention to a new spatial location (Muller and Rabbitt, 1989).

## Acknowledgments

This work was supported by National Institutes of Health Grant NIH MH095084 to B.R.P.

## Citation Gender Diversity Statement

A retrospective analysis of citation patterns in five broad scope neuroscience journals over the past 25 years has revealed a persistent pattern of gender imbalance: Although the proportions of authorship teams (categorized by estimated gender identification of first author/last author) publishing in these journals in 2018 were M(an)/M = .50, W(oman)/M = .27, M/W = .13, and W/W = .10, the comparable proportions for the articles that these authorship teams cited were M/M = .617, W/M = .236, M/W = .09, and W/W = .058 (Dworkin et al., 2020; https://doi.org/10.1038/s41593-020-0658-y). Using a similar method, we estimate the citations by gender category of this paper to be: M/M = .818, W/M = .061, M/W = .091, and W/W = .03

